# Kinetics and functional consequences of BK Channels activation by N-type Ca^2+^ channels in the dendrite of mouse neocortical layer-5 pyramidal neurons

**DOI:** 10.1101/2023.10.26.564136

**Authors:** Laila Ananda Blömer, Elisabetta Giacalone, Fatima Abbas, Luiza Filipis, Domenico Tegolo, Michele Migliore, Marco Canepari

## Abstract

The back-propagation of an action potential (AP) from the axon/soma to the dendrites plays a central role in dendritic integration. This process involves an intricate orchestration of various ion channels, but a comprehensive understanding of the contribution of each channel type remains elusive. In this study, we leverage ultrafast membrane potential recordings (V_m_) and Ca^2+^ imaging techniques to shed light on the involvement of N-type voltage-gated Ca^2+^ channels (VGCCs) in layer-5 neocortical pyramidal neurons’ apical dendrites. We found a selective interaction between N-type VGCCs and large-conductance Ca^2+^-activated K+ channels (BK CAKCs). Remarkably, we observe that BK CAKCs are activated within a mere 500 µs after the AP peak, preceding the peak of the Ca^2+^ current triggered by the AP. Consequently, when N-type VGCCs are inhibited, the early broadening of the AP shape amplifies the activity of other VGCCs, leading to an augmented total Ca^2+^ influx. A NEURON model, constructed to replicate and support these experimental results, reveals the critical coupling between N-type and BK channels. This study not only redefines the conventional role of N-type VGCCs as primarily involved in presynaptic neurotransmitter release but also establishes their distinct and essential function as activators of BK CAKCs in neuronal dendrites. Furthermore, our results provide original functional validation of a physical interaction between Ca^2+^ and K^+^ channels, elucidated through ultrafast kinetic reconstruction. This insight enhances our understanding of the intricate mechanisms governing neuronal signaling and may have far-reaching implications in the field.

## Introduction

In neocortical pyramidal neurons, action potentials (APs) generated in the axon initial segment actively propagate back into the dendritic tree (Stuart and Sakmann, 1994; Stuart et al., 1997), where they elicit transient elevations of intracellular Ca^2+^ concentration (Markram et al., 1995; Schiller et al., 1995). Pharmacological analyses performed in dissociated pyramidal neurons (Stewart and Foehring, 2000) and in Layer-5 (L5) pyramidal neurons from brain slices (Markram et al., 1995; Almog and Korngreen, 2009) have shown that all high-voltage activated (HVA) VGCCs, namely L-type (Ca_v_1), P/Q-type (Ca_v_2.1), N-type (Ca_v_2.2) and R-type (Ca_v_2.3), contribute to the AP-mediated dendritic Ca^2+^ transient, but low-voltage activated (LVA) VGCCs (T-type, Ca_v_3) may also contribute since the expression of these channels in this neuronal type has been reported (Talley et al., 1999). Following the Ca^2+^ transient, Ca^2+^-binding proteins can be activated by indistinct cytosolic Ca^2+^ elevation (Ghosh and Greenberg, 1995), and in this case these proteins can be equally activated by Ca^2+^ ions from any source contributing to the signal. Alternatively, Ca^2+^-binding proteins can be more selectively activated by a physically-coupled Ca^2+^ source. In this case, the protein experiences a larger Ca^2+^ elevation in a restricted nanodomain adjacent to the specific Ca^2+^ source, whereas Ca^2+^ ions from others sources would be less effective. This type of multi-protein structure is well characterized in the synaptic cleft (Gandini and Zamponi, 2022).

VGCCs in pyramidal neuron dendrites target, among other proteins, Ca^2+^-activated K^+^ channels (CAKCs), in particular SK and BK CAKC (Sah and Davies, 2000). Both channels can be physically coupled with a Ca^2+^ source (Vierra and Trimmer, 2022), but it was suggested that BK CAKCs must be localized closer to the Ca^2+^ source to ensure reliable Ca^2+^-dependent activation because they have lower affinity for Ca^2+^ with respect to SK CAKCs (Fakler and Adelman, 2008; Shah et al., 2022). In cell-attached patches from isolated CA1 hippocampal pyramidal neurons it was shown that L-type VGCCs activate exclusively SK CAKCs whereas N-type VGCCs activate exclusively BK CAKCs (Marrion and Tavalin, 1998). In contrast, whole-cell patch-clamp recordings from freshly dissociated neocortical pyramidal neurons showed that BK CAKCs are activated both by L-type and N-type VGCCs in this preparation (Sun et al., 2003), consistently with the findings that both L-type VGCCs (Grunnet and Kaufmann, 2004) and N-type VGCCs (Loane et al., 2007) can molecularly co-assemble with BK CAKCs. Finally, it was reported that SK channels are selectively activated by R-type VGCCs in basal dendrites adjacent to dendritic spines (Jones and Stuart, 2013). From the functional point of view, CAKCs participate in shaping the AP, but the kinetics of activation is different for SK and BK CAKCs (Sah and Faber, 2000) as also reproduced by computer modeling (Almog and Korngreen, 2014). Specifically, SK CAKCs regulate the AP waveform during the medium and late phase of the AP re-polarisation with a variability that depends on the different Ca^2+^ source and neuronal types (Pedarzani and Stocker, 2004; Bond et al., 2004; Villalobos et al., 2004). Interestingly, dendritic excitability is impacted by SK CAKC activation in unpredictable manner (Bock et al., 2019). In contrast, BK CAKCs, which are functionally expressed in the dendrites L5 neocortical pyramidal neurons (Kang et al., 1998; Benhassine and Berger, 2005) regulate the AP in the early phase of re-polarisation (Sun et al., 2003) when the membrane potential (V_m_) is depolarized, consistently with the voltage dependence of the channel (Vergara et al., 1998; Cui, 2010).

The close interaction and selective coupling between the Ca^2+^ source and the BK CAKC implies the activation of the K^+^ channel at sub-millisecond time scale (Berkfeld et al., 2006). In the present study, performed in L5 pyramidal neurons of the somatosensory cortex, we used ultrafast optical measurements of V_m_ (Popovic et al., 2015) and Ca^2+^ currents (Jaafari et al., 2014) to investigate in parallel the kinetics of the dendritic back-propagating AP (bAP) and that of the associated Ca^2+^ current. We found that BK CAKCs are selectively activated by N-type VGCCs. Yet, whereas the peak of the Ca^2+^ current is delayed by >500 µs with respect to the bAP peak, consistently with the kinetics of activation and deactivation of VGCCs in the order of 1 ms (Kay and Wong, 1987), the activation of BK CAKCs occurs in the first 500 µs, providing a negative feedback to the cytosolic Ca^2+^ elevation (Shah et al., 2022). We built a realistic NEURON model showing that experimental results could be reproduced by the only when N-type VGCCs and BK CAKCs were physically interacting. As a consequence, we demonstrate that combined ultrafast V_m_ and Ca^2+^ imaging can provide kinetic evidence of physical interactions between a Ca^2+^ source and a Ca^2+^-binding partner.

## Materials and Methods

### Brain slices preparation and maintenance

All experiments were performed in the “Laboratory of Interdisciplinary Physics” and mice manipulations previous to euthanasia were done in accordance with European Directives 2010/63/UE on the care, welfare and treatment of animals. Specifically, procedures were reviewed by the ethics committee affiliated to the animal facility of the university (D3842110001). Mice (C57BL/6j, 21-35 postnatal days old) purchased from Janvier Labs (Le Genest-Saint-Isle, France) were anesthetised by isofluorane inhalation and decapitated to extract the entire brain. Neocortical slices (350 µm thick) were prepared as described in recent previous reports (Filipis and Canepari, 2021; Montnach et al., 2022; Filipis et al., 2023), using a Leica VT1200 vibratome (Wetzlar, Germany). The extracellular solution contained (in mM): 125 NaCl, 26 NaHCO_3_, 1 MgSO_4_, 3 KCl, 1 NaH_2_PO_4_, 2 CaCl_2_ and 20 glucose, bubbled with 95% O_2_ and 5% CO_2_. Slices were incubated at 37°C for 45 minutes and maintained at room temperature before being transferred to the recording chamber where the temperature was maintained at 32-34°C.

### Electrophysiology and imaging

Slices with L5 pyramidal neurons in the somato-sensory cortex having the initial part of the apical dendrite parallel to the surface were used for the experiments. Patch-clamp (whole-cell) recordings were made using a Multiclamp 700A (Molecular Devices, Sannyvale, CA) with a basic intracellular solution containing (in mM): 125 KMeSO_4_, 5 KCl, 8 MgSO_4_, 5 Na_2_-ATP, 0.3 Tris-GTP, 12 Tris-Phosphocreatine, 20 HEPES, adjusted to pH 7.35 with KOH. To this basic solution, one or more indicators were added according to the type of optical recording. In V_m_ imaging experiments without concomitant Ca^2+^ recordings, cell membranes were loaded with the voltage-sensitive dye D-2-ANEPEQ (JPW1114, 0.2 mg/mL, Thermo Fisher Scientific) for 30 minutes using a first patch clamp recording and then re-patched a second time with dye free solution. In V_m_ imaging experiments with concomitant Ca^2+^ recordings, the intracellular solution in the second patch contained a Ca^2+^ indicator at the concentration of 2 mM which was either Cal-520FF (AAT Bioquest, Sunnyvale, CA, USA, see Blömer et al., 2021) in 2 cells or Fura2FF (Santa Cruz Biotechnology) in 6 cells. The choice of using, in some experiments, a UV-excitable indicator is motivated by the need to obtain, in the majority of cells, both V_m_ and Ca^2+^ signals with a combination of spectrally non-overlapping indicators (Vogt et al., 2011a). Finally, in exclusive Ca^2+^ imaging experiments, the Ca^2+^ indicator added to the intracellular solution at the concentration of 2 mM was either Oregon Green BAPTA-5N (Thermo Fisher Scientific) or the most sensitive low-affinity dye Cal-520FF. Ca^2+^ recordings started 30 minutes after achieving the whole cell configuration, a time necessary for full dye equilibration in the initial 200 µm segment of the apical dendrite. Somatic APs were elicited by short current pulses of 1.5-2.5 nA through the patch pipette in current clamp mode. Electrical somatic V_m_ transients were acquired at 20 kHz and the bridge associated with the current pulse was corrected offline by using the recorded injected current. Finally, the measured V_m_ was corrected for −11 mV junction potential. Experiments were performed using an imaging system described in several previous reports (Jaafari et al., 2014; Jaafari and Canepari, 2016; Ait Ouares et al. 2019; Ait Ouares and Canepari 2020). This system is based on an Olympus BX51 microscope equipped with a 60X/1.0 NA Nikon objective where whole-field fluorescence illumination is provided either with an OPTOLED system (Cairn Research, Faversham, UK) or with an LDI-7 laser (89 North, Williston, VT). Fluorescence emission was demagnified by 0.5X and acquired with a NeuroCCD camera (Redshirt Imaging, Decatur, GA). Signals were sampled at 20 kHz (for 8 ms) with a resolution of 4X26 pixels except in V_m_ imaging calibration experiments where signals were sampled at 5 kHz with a resolution of 26X26 pixels (for 160 ms). In V_m_ imaging recordings, JPW1114 fluorescence was excited using the 528 nm line of an LDI-7 laser (89 North, Williston, VT) and the emitted fluorescence was long-pass filtered at >610 nm before being acquired. In Ca^2+^ imaging recordings, OG5N or Cal-520FF were excited by the 470 nm line of the OPTOLED and the emitted fluorescence was band-pass filtered at 530 ± 21 nm before being acquired. Finally, in Ca^2+^ imaging recordings with Fura2FF, fluorescence was excited by the 385 nm line of the OPTOLED and the emitted fluorescence was band-pass filtered at 510 ± 41 nm before being acquired.

### Pharmacology of Ca^2+^ and K^+^ channels

Channels blockers used in this study were either peptides (animal toxins), purchased from Smartox Biotechnology (Saint Egrève, France), or smaller organic molecules. Peptides, dissolved in water and used at the final concentration of 1 µM, were: ω-agatoxin-IVA (P/Q-type VGCC blocker), ω-conotoxin-GVIA (N-type VGCC blocker), snx-482 (R-type VGCC blocker), apamin (SK CAKC blocker) and iberiotoxin (BK CAKC blocker). The blockade of L-type VGCCs was achieved using 4-(2,1,3-Benzoxadiazol-4-yl)-1,4-dihydro-2,6-dimethyl-3,5-pyridinecarboxylic acid methyl 1-methylethyl ester (isradipine, purchased from HelloBio, Bristol, UK) that was diluted in external solution at 20 µM concentration. To obtain an alternative blockade of N-type VGCCs we used *N*-[[4-(1,1-Dimethylethyl)phenyl]methyl-*N*-methyl-L-leucyl-*N*-(1,1-dimethylethyl)-*O*-phenylmethyl)-L-tyrosinamide (pd173212, purchased from TOCRIS, Bristol, UK) that was used at the final concentration of 5 µM. Finally, to block T-type VGCCs, we used a cocktail of two drugs (from TOCRIS) at the final concentrations of 30 µM and 5 µM respectively: 3,5-dichloro-*N*-[[(1α,5α,6-exo,6α)-3-(3,3-dimethylbutyl)-3-azabicyclo[3.1.0]hex-6-yl]methyl]-benzamide-hydrochloride (ml218) and (1*S*,2*S*)-2-[2-[[3-(1*H*-Benzimidazol-2-yl)propyl]methylamino]ethyl]-6-fluoro-1,2,3,4-tetrahydro-1-(1-methylethyl)-2-naphthalenyl-cyclopropanecarboxylate-dihydrochloride (nnc550396). In all experiments, the blocker(s) was (were) locally delivered by gentle pressure application near the optical recording area using a pipette of 2-4 µm of diameter. The application lasted 2-4 minutes before recording in order to assure the stable blockade of the channel(s).

### Data analysis and quantification

All data were from averages of 6-9 trials with identical somatic response in Ca^2+^ recordings, and of 4 trials with identical somatic response in V_m_ recordings. Raw data were converted in MATLAB format and analysed using custom-made code written in MATLAB. As a first step, fluorescence values including an AP were corrected for photo-bleaching using multi-exponential fits of fluorescence values in single trials without an AP. Then, the fractional change of fluorescence from the first image (ΔF/F_0_) was calculated over the mean fluorescence in regions of 20-40 µm located at ∼100 µm from the soma. It is important to state that all the conclusions reported in this study were based on comparisons of signals under control conditions and signals after the blockade of one or more channels, for which a calibration of the size of the signals was not necessary. Nevertheless, to build realistic NEURON models, the V_m_ (ΔF/F_0_) associated with the bAP was converted into mV considering an attenuation of the somatic AP size, which was 4% on average. This value was obtained in a set of experiments in which the V_m_ (ΔF/F_0_) associated with the bAP was calibrated in mV using a previously reported method (Vogt et al., 2011b). As shown in the example of Fig.1*A*, the V_m_ ΔF/F_0_ signal associated with an AP was measured in the apical dendrite at ∼100 µm from the soma. Then, in the presence of 1 µM tetrodotoxin to block APs and 50 µM cyclothiazide to inhibit AMPA receptors desensitization, L-glutamate was locally photo-released from 1 mM 4-Methoxy-7-nitroindolinyl-caged-L-glutamate (MNI-glutamate, TOCRIS) using an OPTOLED pulse of 1 ms at 365 nM wavelength. As this procedure brings the dendritic V_m_ to the reversal potential of ionotropic glutamate receptors (i.e. to 0 mV), the associated V_m_ ΔF/F_0_ signal was used to calibrate the V_m_ ΔF/F_0_ signal associated with the bAP. This assessment was repeated in N = 8 cells obtaining a consistent result (Fig.1*A*). For the quantification of Ca^2+^ signals, the dendritic Ca^2+^ ΔF/F_0_ signal associated with the bAP, normalized to its asymptotic value, was initially fitted with a 4-sigmoid function Y(t):

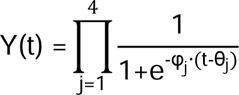

where t is time and φ_j_ and θ_j_ are the parameters to be determined by the fit. The time derivative of Y(t) was then used as measurement of the Ca^2+^ current (I_Ca_). The maxima of Y(t) and of its derivative were used to quantify the fractional change of the signals produced by the channel blockers.

**Figure 1.**
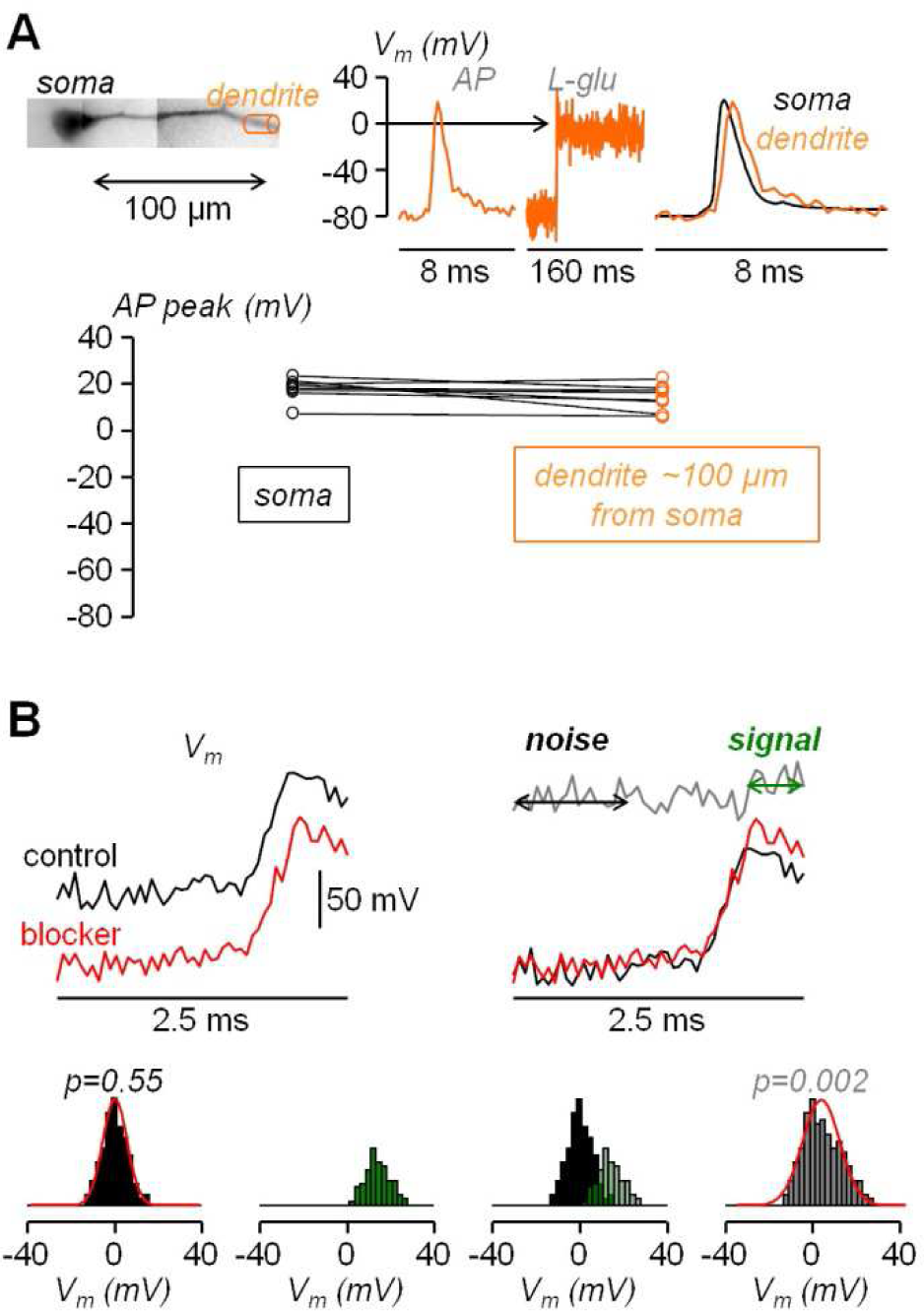
Calibration of dendritic AP and statistical assessment of the AP widening following pharmacological manipulation. ***A***, Top-left, reconstruction of L5 pyramidal neuron with dendritic compartment at ∼100 µm distance from the soma outlined by a cylinder (orange). Top-right, V_m_ optical signal associated with an AP in the region on the left, calibrated using a depolarization to 0 mV obtained by uncaging L-glutamate (see Materials and Methods), showing a slight attenuation of the dendritic bAP with respect to the somatic AP (black trace). Bottom, AP peak in the soma and in the dendrite at ∼100 µm distance from the soma measured in N = 8 cells. ***B***, Top, to simulate an experimental widening two sets of normally distributed random values with 5 mV standard deviation are added to a somatic AP recorded at 20 kHz, once in control conditions (black traces) and once in the presence of the channel blocker (red traces); the normal distributions are centered at 0 mV except for the 10 points (500 µs) following the AP peak in the red trace for which the distribution is centered at +15 mV; the gray trace on the right shows the difference between the red and black trace; the analysis is performed over the 20 points (1 ms) preceding the current injection (noise) and the 10 points following the AP peak (signal). Bottom, following the procedure illustrated above applied to N = 8 cells to obtain 160 points for the noise and 80 points for the signal, histograms of noise (black columns), of the signal (green columns) and of the noise+signal (240 points, gray columns); to qualitatively visualize the consistency with normality, normal distributions with mean and standard deviation calculated from noise and from the noise+signal are superimposed to the respective histograms (red traces); the p values of the two Lilliefors tests are indicated.

### Statistics

To assess the consistency of the experimental results obtained in groups of cells, three statistical tests were performed and in all tests we considered 0.01 as the threshold for significance.

1. To assess the effect of a channel blocker on the Ca^2+^ and I_Ca_ signals in the same group of experiments, the parametric paired t-test was performed on the signals maxima under two different conditions. In each cell, the maxima were measured in control conditions and in the presence of one or several channel blockers.
2. To assess the effect of a channel blocker on the Ca^2+^ and I_Ca_ signals in two different groups of experiments, the Wilcoxon rank non-parametric test was performed on the fractional changes of signals maxima under two different conditions in the two groups of cells.
3. Finally, to establish the effect of a channel blocker on the shape of the dendritic AP, we assessed the widening of the AP over the photon noise of the recording using the following statistical analysis. Assuming that the photon noise is normally distributed, the distribution of the difference between the noise samples in the presence of the channel blocker and in control condition is also normal with standard deviation equal to √2 times the standard deviation of the noise. Thus the hypothesis to test was whether or not the addition of the sample differences at given intervals after the AP peak produces a distribution that deviates from normality. The rationale of the analysis is illustrated in the simulations reported in Fig.1*B*. In the example reported on the top of the panel, two sets of normally distributed values with 5 mV standard deviation (∼15 mV peak-to-peak noise) were added to a somatic AP recorded at 20 kHz. This was done once to simulate the control conditions and once to simulate the presence of the channel blocker. The normal distributions were centered at 0 mV except for the 500 µs following the AP peak in the red trace for which the distribution was centered at +15 mV in order to mimic the widening. The distribution of the difference between the noise samples in the presence of the channel blocker and in control condition are also normal with standard deviation equal to √2 times the standard deviation of the original signals. The distribution of the difference between the noise + signal samples, however, deviates from normality since the two original distributions are centered at two different values. Thus, we repeated the simulations in N = 8 cells and we illustrate the histograms of the difference in the noise, in the signal and in the noise + signal (Fig.1*B*). To visualize the normal behavior of the noise difference and the deviation from normality of the noise + signal difference, we superimpose to the histograms the two normal distributions with mean and standard deviation calculated from the points. The test to obtain a quantitative assessment of the normal behavior is, in principle the Kolmogorov-Smirnov test. In practice, since in the experiments the noise varies from one cell to another, we opted for the stronger Lilliefors test, which is a generalization of the Kolmogorov-Smirnov test for unknown normal distributions (Lilliefors, 1967). The p values of this test for the above simulations are reported in Fig.1*B*. In the experiments, to establish the kinetics of the widening, the test was repeated over three distinct time intervals during the falling phase of the AP, namely the first 500 µs following the AP peak, the next 500 µs and the next 1.5 ms.

### Computational modelling

All simulations were carried out using the NEURON simulation environment (v.8.2.0). Model and simulation files will be available in the ModelDB section of the Senselab database (https://modeldb.science/2015410). We used a 3D reconstructed morphology of a mouse L5 pyramidal neuron (morphology NMO_36595, Buchanan et al., 2012) downloaded from NeuroMorpho.org database (Ascoli et al., 2007), with uniform passive properties (Cm = 1 mF/cm^2^, Rm = 30 kohm/cm^2^, Ra = 160 ohm·cm) where we attached the axon morphology used in Filipis et al. (2023). Temperature was set at 33°C. The NEURON model, built based on the original model reported by Hallermann et al. (2012) was previously employed in our previous studies (Filipis & Canepari, 2021, Filipis et al., 2023). It was adapted to reproduce the somatic AP and dendritic bAP recordings used as a reference in this work. Active properties included two Na^+^ currents, the Na_v_1.2 and Na_v_1.6 channels, five types of K^+^ currents (a delayed rectifier conductance, A-type K^+^ conductance, K_v_7 conductance, SK Ca^2+^-dependent K^+^ conductance and BK SK Ca^2+^-dependent K^+^ conductance), a non-specific I_h_ current, Ca^2+^ conductance modelling including T-type, L-type, R-type, N-type and P/Q-type Ca^2+^ currents, and a simple Ca^2+^-extrusion mechanism with a 20 ms time constant, to reproduce the measured Ca^2+^ transient decay. For T-type VGCCs, we used the model from Migliore et al. (2008). For R-type VGCCs, we used the model from Mandge and Manchanda (2018). For N-type, P/Q-type, and L-type VGCCs, we used the same model from Wimmer et al. (2010). For BK channel in dendrites, we modified the model in Filipis et al. (2023) to match experimental data on the bAP. In particular, we shifted the voltage dependent activation kinetic by +10mV and used a time constant with a sigmoidal form. All other channels were taken from Filipis et al. (2023). Relative spatial distributions of channel dendritic densities were in accordance with Migliore and Shepherd (2002) and Ramaswamy and Markram (2015); Na_v_1.2 expression was ∼50 fold lower compared to the axon initial segment (Kole et al., 2008); an inward current generated by HCN cation channels was also inserted with a density increasing exponentially to 50 fold in the distal apical dendrite compared to the soma (Harnett et al., 2015). To determine the absolute channel densities of the model, we initially used the calibrated voltage imaging recording in Fig.1A to reflect the action potential experimental traces in both the soma and the dendrite. Specifically, we first modified the Na^+^ channels to match the somatic AP and we then modified Ca^2+^ and K^+^ channels to match the dendritic bAP. After that, we refined the dendritic channel density for each individual type of VGCC and set a coupling between particular VGCCs and the BK CAKC. To mimic this coupling, we distinguished the components of Ca^2+^ influx from each individual VGCC and we introduced an affinity of the CAKC for a specific component multiplied a factor α > 0 (α = 0 means no coupling). In agreement with a study in neocortical pyramidal neurons (Sun et al., 2003), we established the coupling both for L-type and N-type VGCCs, using values of α = 3 and α = 10 to mimic “weak” and “strong” coupling respectively. Finally, the pharmacological blockade of each channel in the experiments was mimicked by removing 90% of the channel density from the model.

### Data and code availability

The complete dataset of imaging/electrophysiological experiments in brain slices, used in this study, is available in the public repository Zenodo (doi: 10.5281/zenodo.7623898). The Neuron model is available in the NeuronDB database at: https://modeldb.science/2015410. Matlab codes used for data analysis are available at https://github.com/MarcoCanepari/NAV12-BK-paper.

## Results

### The dendritic Ca^2+^ influx associated with the bAP is mediated by diverse VGCCs

This study started with the investigation of the VGCCs that mediate the Ca^2+^ influx associated with the bAP in the apical dendrite at ∼100 µm from the soma using the same approach already utilised in CA1 hippocampal pyramidal neuron (Jaafari and Canepari, 2016). In the present experiments, L5 pyramidal neurons were loaded with 2 mM of a low-affinity Ca^2+^ indicator, either OG5N or the more sensitive Cal-520FF (Blömer et al. 2021), and Ca^2+^ transients (*ΔF/F_0_* signals) were measured at 20 kHz by averaging fluorescence from dendritic segments of 20-40 µM length at ∼100 µm from the soma. To investigate the contribution of the diverse VGCCs, we locally delivered one or several channel blockers using a pipette positioned near the recording region, as already done in other studies (Ait Ouares et al., 2019; Filipis et al., 2023). Specifically, we blocked L-type VGCCs with 20 µM isradipine, P/Q-type VGCCs with 1 µM ω-agatoxin-IVA, N-type VGCCs with 1 µM ω-conotoxin-GVIA, R-type VGCCs with 1 µM of SNX482, and T-type VGCCs with 5 µM ML218 and 30 µM NNC550396. Fig. 2*A* shows a Ca^2+^ transient associated with a bAP that was fully inhibited by the cocktail of all VGCC blockers, a result consistently observed in 6 cells tested. Fig. 2*A* also shows the analysis of the Ca^2+^ transient consisting in fitting the signal with a 4-sigmoid function and to calculate the time-derivative of the fit to extrapolate the kinetics of the Ca^2+^ current (I_Ca_). We applied this analysis on the Ca^2+^ transient recorded first in control solution and then after locally blocking each individual VGCC. The five representative examples reported Fig. 2*B* show that the blockade of L-type, P/Q-type, R-type and T-type VGCCs decreased, at different extent, the size of both ΔF/F_0_ and I_Ca_ signals, but surprisingly the blockade of N-type VGCCs increased the size of both ΔF/F_0_ and I_Ca_ signals. The test was repeated in 7-10 different cells for each channel blocker and the effect of the inhibitor was quantified by measuring the maximum of the ΔF/F_0_ and I_Ca_ signals (Fig. 2*C*). Except for the blockade of N-type VGCCs, the individual blockade of each VGCC type produced on average a decrease of both *ΔF/F_0_* and *I_Ca_*signals, an effect that was more important for L-type and T-type VGCCs. In the case of N-type VGCCs, the blockade increased both ΔF/F_0_ and I_Ca_ signals in 8/11 cells tested. The unexpected result of locally delivering 1 µM ω-conotoxin-GVIA could be explained by the coupling of the N-type VGCC with a mechanism that boosts the Ca^2+^ influx associated with the bAP, or with the toxin binding to a different target. To discriminate between these two hypotheses, we repeated the analysis of the Ca^2+^ transient of Fig. 2 by blocking the N-type VGCC with another selective inhibitor, namely pd173212 (Hu et al., 1999). As shown in the two representative examples of Fig. 3*A-B*, local delivery of either 5 µM pd173212 or 1 µM ω-conotoxin-GVIA increased the size of both ΔF/F_0_ and I_Ca_ signals in a qualitatively similar manner. Altogether, the *ΔF/F_0_* signal increased in 7 cells tested after delivery of pd173212, and the I_Ca_ signal increased in 4/7 cells tested after delivery of pd173212 (Fig. 3*C*). We concluded that the blockade N-type VGCCs boosts the Ca^2+^ influx associated with the bAP.

**Figure 2.**
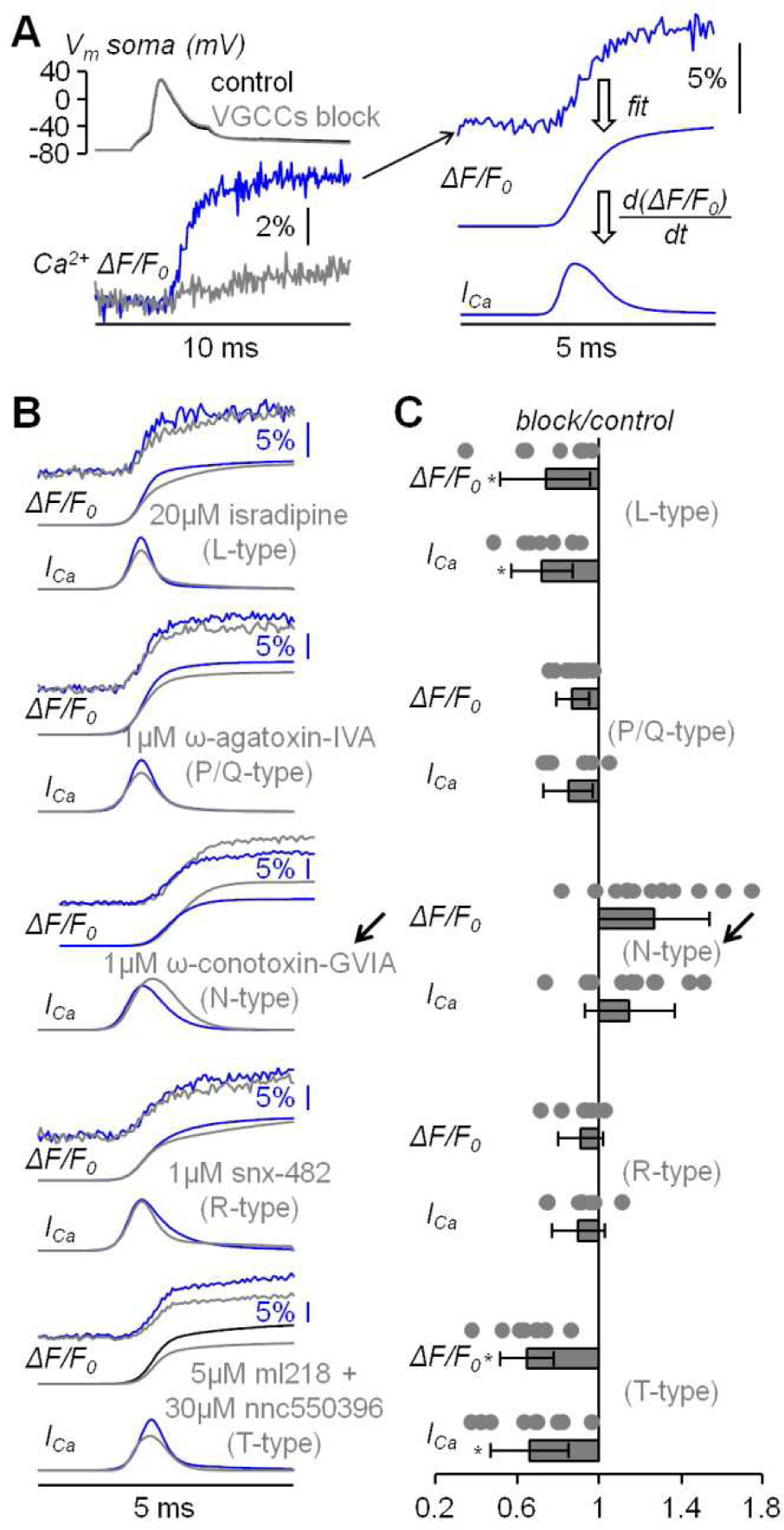
Analysis of the diverse VGCCs mediating the Ca^2+^ transient associated with the bAP. ***A***, Left V_m_ somatic recording (top) and Ca^2+^ transient (bottom) in control solution and after blocking all VGCCs. Right, procedure of analysis of the raw Ca^2+^ transient (top trace) with a 4-sigmoid fit (middle trace) and the calculation of the time-derivative (*I_Ca_*, bottom trace). ***B***, Five representative examples of Ca^2+^ transients with 4-sigmoid fit (*ΔF/F_0_*, top traces) and time derivative (*I_Ca_*, bottom traces) in control solution and after local delivery of either 20 µM of the L-type VGCC inhibitor isradipine, 1 µM of the P/Q-type VGCC inhibitor ω-agatoxin-IVA, 1 µM of the N-type VGCC inhibitor ω-conotoxin-GVIA, 1 µM of the R-type VGCC inhibitor snx-482 or 5 µM and 30 µM, respectively, of the T-type VGCC inhibitors ml218 and nnc550396. The arrow indicates the unexpected boosting of the Ca^2+^ transient produced by ω-conotoxin-GVIA. ***C***, Single values and fractional change with respect to control of the maxima of the *ΔF/F_0_*and *I_Ca_* signals for the blockade of L-type VGCCs (N = 7, *ΔF/F_0_*: 0.74 ± 0.22, *I_Ca_*: 0.72±0.15); for the blockade of P/Q-type VGCCs (N = 8, *ΔF/F_0_*: 0.87 ± 0.08, *I_Ca_*: 0.85 ± 0.12); for the blockade of N-type VGCCs (N = 11, *ΔF/F_0_*: 1.27 ± 0.27, *I_Ca_*: 1.15 ± 0.22); for the blockade of R-type VGCCs (N = 7, *ΔF/F_0_*: 0.91 ± 0.11, *I_Ca_*: 0.90 ± 0.13); for the blockade of T-type VGCCs (N = 7, *ΔF/F_0_*: 0.65 ± 0.13, *I_Ca_*: 0.66 ± 0.19). “*” indicates that p < 0.01 in paired t-test performed on the signal maxima in control solution and after delivery of the channel blocker.

**Figure 3.**
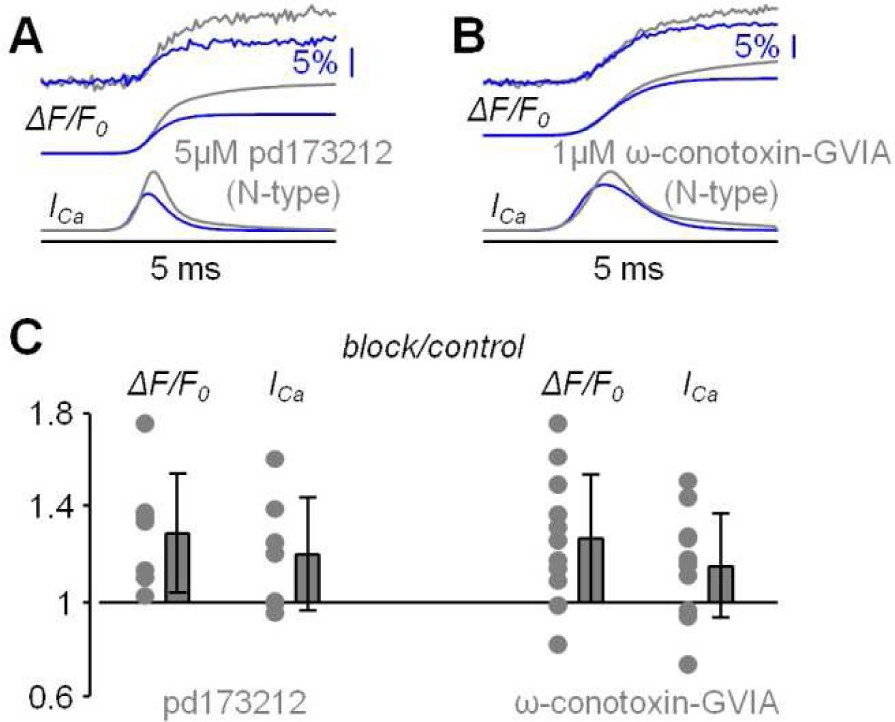
Analysis of the blockade of N-type VGCCs using pd173212. ***A***, Representative example of Ca^2+^ transient with 4-sigmoid fit (*ΔF/F_0_*, top traces) and time derivative (*I_Ca_*, bottom traces) in control solution and after local delivery of 5 µM of the N-type VGCC inhibitor pd173212. ***B***, Another representative example of Ca^2+^ transient with 4-sigmoid fit (*ΔF/F_0_*, top traces) and time derivative (*I_Ca_*, bottom traces) in control solution and after local delivery of 1 µM of the N-type VGCC inhibitor ω-conotoxin-GVIA. ***C***, Left, single values and fractional change with respect to control of the maxima of the *ΔF/F_0_* and *I_Ca_* signals for the blockade of N-type VGCCs with pd173212 (N = 7, *ΔF/F_0_*: 1.29 ± 0.25, *I_Ca_*: 1.20 ± 0.24). The single values and fractional change with respect to control of the maxima for the blockade of N-type VGCCs with ω-conotoxin-GVIA are reported on the right for comparison.

### N-type VGCCs are coupled with BK CAKCs

The boosting of the Ca^2+^ influx observed in Fig. 2 and Fig. 3 can be due to an enhancement of the depolarisation produced by the loss of Ca^2+^ influx *via* N-type VGCCs. It was shown in freshly dissociated neocortical pyramidal neurons of the mouse that BK CAKCs are activated by N-type VGCCs (Sun et al., 2003). Thus, we repeated the analysis of the Ca^2+^ transient of Fig. 2 and Fig. 3 by blocking in sequence first BK CAKCs with local delivery of 1 µM iberiotoxin and then BK CAKCs and N-type VGCCs together with local delivery of 1 µM iberiotoxin and 1 µM ω-conotoxin-GVIA. In the representative example of Fig. 4*A*, the blockade of BK CAKCs increased both ΔF/F_0_ and I_Ca_ signals, but the further blockade of N-type VGCCs decreased both ΔF/F_0_ and I_Ca_ signals. This result was consistently obtained in N = 9 cells tested. As shown in Fig. 4*B*, the blockade of BK CAKCs boosted the Ca^2+^ transient similarly to the blockade of N-type VGCCs. In contrast, the blockade of N-type VGCCs in the presence of iberiotoxin reduced the Ca^2+^ transient. To statistically assess the consistency of this result we performed a Wilcoxon rank non-parametric test on the groups of cells where a blocker was applied under different conditions, using p<0.01 as discriminator to establish whether two groups were different. As show in Table 1, the effects on both ΔF/F_0_ and I_Ca_ signals were similar when comparing the groups of cells where ω-conotoxin-GVIA, pd173212 or iberiotoxin were applied in control solutions, or when the effects of iberiotoxin were compared with the ensemble of cells where N-type VGCCs were blocked in control solution. In contrast, the effects on both ΔF/F_0_ and I_Ca_ signals were different when comparing the cells where ω-conotoxin-GVIA was applied in the presence of iberiotoxin with the groups of cells where ω-conotoxin-GVIA or pd173212 were applied in control solution, or when the comparison was done with the ensemble of cells where N-type VGCCs were blocked in control solution. Since the blockade of BK CAKCs prevents the boosting of the Ca^2+^ influx when blocking N-type VGCCs, we concluded that this effect was due to the functional coupling of the two channels.

**Figure 4.**
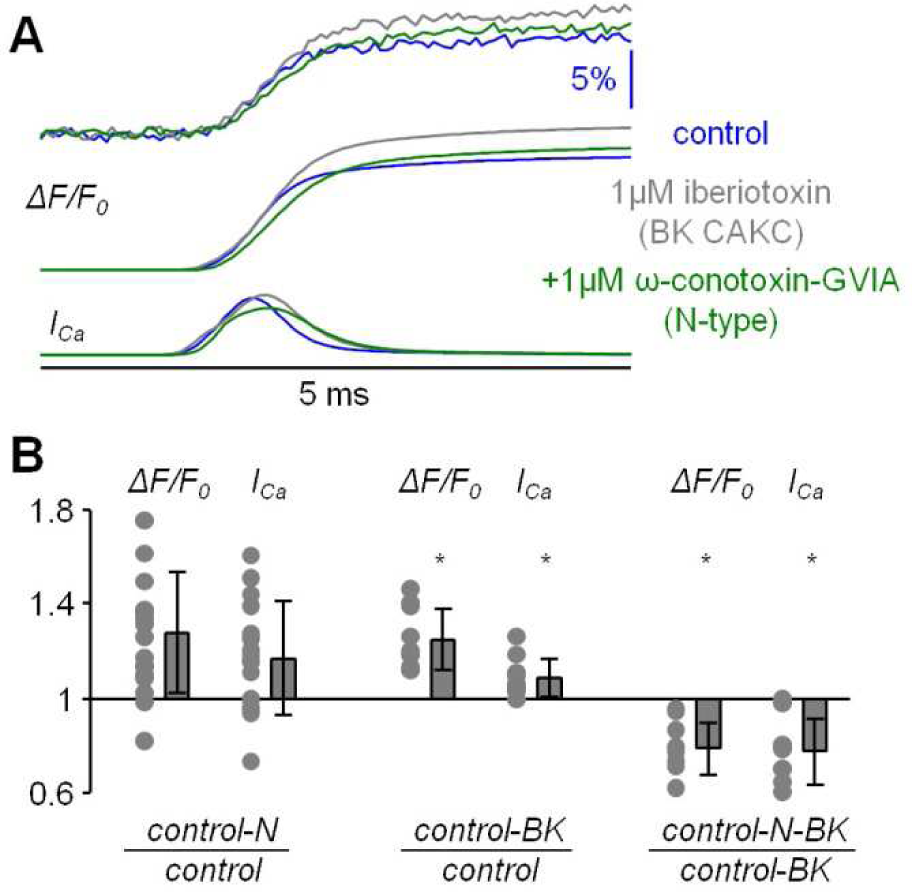
Analysis of the blockade of N-type VGCCs after blocking BK CAKCs. ***A***, Representative example of Ca^2+^ transient with 4-sigmoid fit (*ΔF/F_0_*, top traces) and time derivative (*I_Ca_*, bottom traces) in control solution, after local delivery of 1 µM of the BK CAKCs inhibitor iberiotoxin and after further delivery of 1 µM of the N-type VGCC inhibitor ω-conotoxin-GVIA. ***B***, Left, single values and fractional change with respect to control solution of the maxima of the *ΔF/F_0_* and *I_Ca_* signals for the blockade of N-type VGCCs (either with ω-conotoxin-GVIA, N = 18, *ΔF/F_0_*: 1.27 ± 0.26, *I_Ca_*: 1.17 ± 0.22). Center, single values and fractional change with respect to control solution of the maxima of the *ΔF/F_0_*and *I_Ca_* signals for the blockade of BK CAKCs (N = 9, *ΔF/F_0_*: 1.25 ± 0.13, *I_Ca_*: 1.09 ± 0.08). Right, single values and fractional change with respect to the blockade of BK CAKCs only of the maxima of the *ΔF/F_0_* and *I_Ca_* signals for the combined blockade of N-type VGCCs and BK CAKCs (with ω-conotoxin-GVIA, N = 9, *ΔF/F_0_*: 0.79 ± 0.11, *I_Ca_*: 0.78 ± 0.14). “*” indicates that p < 0.01 in paired t-test performed on the signal maxima.

**Table 1.** Wilcoxon rank non-parametric test performed on the effects on Ca^2+^ transients produced by blocking N-type VGCCs and BK CAKCs. The considered groups are: delivery of 1 µM ω-conotoxin-GVIA in control conditions (N = 11 cells); delivery of 5 µM pd173212 in control conditions (N = 7 cells); delivery of either 1 µM ω-conotoxin-GVIA or 5 µM pd173212 in control conditions (N = 18 cells); delivery of 1 µM iberiotoxin in control conditions (N = 9 cells); delivery of 1 µM ω-conotoxin-GVIA in the presence of 1 µM iberiotoxin (N = 9 cells). In each line, the two groups compared are reported in the left columns and the p values for the *ΔF/F_0_* and *I_Ca_* signals are reported on the right columns. Tests where p was < 0.01 are reported in bold characters.

| | | $p \Delta F/F_0$ | $p I_{Ca}$ |
| --- | --- | --- | --- |
| ω-conotoxin-GVIA/control | pd173212/control | 0.8601 | 0.7914 |
| ω-conotoxin-GVIA/control | iberiotoxin/control | 0.9998 | 0.2875 |
| **ω-conotoxin-GVIA/control** | **ω-conotoxin-GVIA/iberiotoxin** | **$4.74 \cdot 10^{-4}$** | **0.0024** |
| pd173212/control | iberiotoxin/control | 0.8371 | 0.7577 |
| **pd173212/control** | **ω-conotoxin-GVIA/iberiotoxin** | **$1.75 \cdot 10^{-4}$** | **0.0012** |
| (ω-conotoxin-GVIA&pd173212)/control | iberiotoxin/control | 0.9385 | 0.3681 |
| **(ω-conotoxin-GVIA&pd173212)/control** | **ω-conotoxin-GVIA/iberiotoxin** | **$6.71 \cdot 10^{-5}$** | **$4.26 \cdot 10^{-4}$** |

### The peak of the Ca^2+^ current is delayed with respect of the bAP peak by >500 µs

Before further analyzing the interaction between N-type VGCCs and BK CAKCs, we compared the kinetics of the dendritic I_Ca_ with that of the eliciting bAP at ∼100 µm from the soma. In a previous study, we showed that in L5 pyramidal neurons the peak of the I_Ca_ occurs before the AP peak in the distal part of the axon initial segment whereas it is delayed in the proximal part (Filipis et al., 2023). In another study, using ultrafast combined V_m_ and Ca^2+^ imaging, we showed that the I_Ca_ was further delayed with respect to the AP peak in the proximal part of the apical dendrite of CA1 hippocampal pyramidal neurons (Jaafari and Canepari, 2016). Thus, we performed the same analysis in the proximal part of the apical dendrite in L5 pyramidal neurons using, as Ca^2+^ indicator, either Cal-520FF or Fura2FF (see Materials and Methods). A representative example of this type of measurement is reported in Fig. 5*A*. In this particular example, the peak of the I_Ca_ corresponding to the maximal slope of the dendritic Ca^2+^ transient is delayed by 650 µs (13 samples) with respect to the dendritic AP. Consistently with the result of this experiment, a delay ranging from 500 µs to 800 µs was observed in N = 8 cells (Fig. 5*A*). From the kinetics of the Ca^2+^ transient we concluded that VGCCs start opening during the rising phase of the AP, but they remain open during the entire falling phase of the AP increasing the cytosolic Ca^2+^ concentration over this period. As a consequence, if the activation of a Ca^2+^-binding protein is linear with the cytosolic Ca^2+^, the maximal effect of this activation must be observed at least 500 µs after the AP peak.

**Figure 5.**
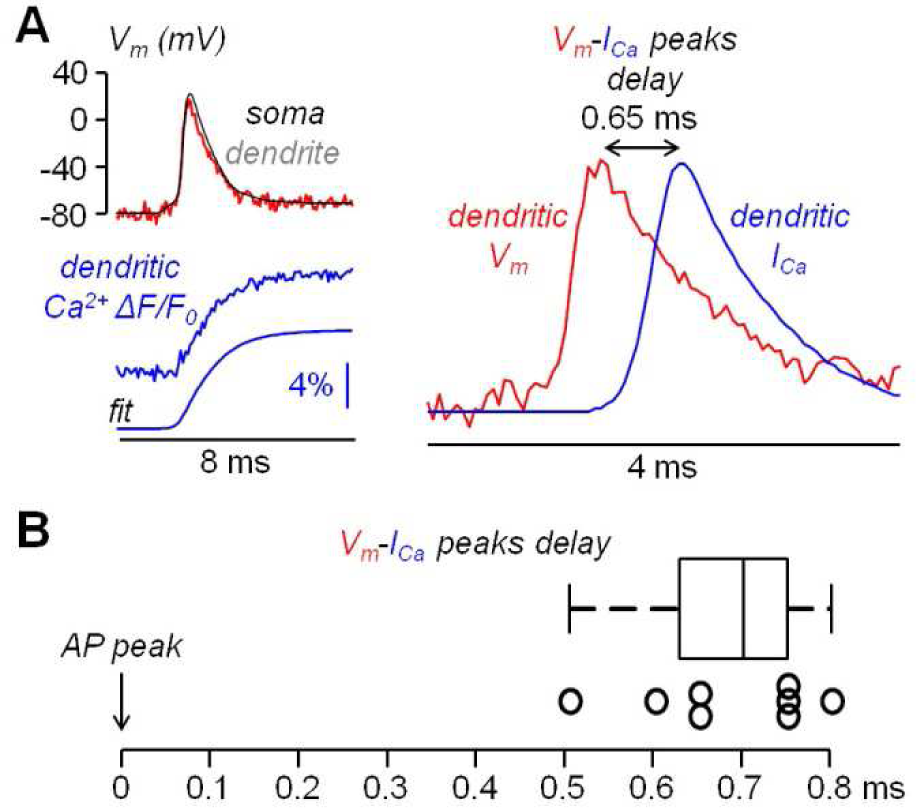
Kinetics of the Ca^2+^ current (I_Ca_) relative to the bAP. ***A***, Left, somatic (black) and dendritic (∼100 µm from the soma, red) AP and concomitant measurement Ca^2+^ transient with 4-sigmoid fit (using Cal-520FF). Right, dendritic AP superimposed to the I_Ca_ (time derivative of the fit) indicating a delay of 0.65 ms between the two signals. ***B***, Delay of the I_Ca_ peak with respect to the AP peak in N = 8 cells tested using combined V_m_ and Ca^2+^ imaging.

### The effect of blocking N-type VGCCs or BK CAKCs occurs in the first 500 µs after the AP peak

If the inhibition of a K^+^ channel occurs during an AP, then this inhibition is expected to change the shape of the AP. Thus, using ultrafast V_m_ imaging (Popovic et al., 2015), as shown in the examples reported in Fig. 6, we investigated the kinetics of the dendritic bAP after blocking L-type VGCCs (panel *A*), P/Q-type VGCCs (panel *B*), N-type VGCCs (panel *C*), R-type VGCCs (panel *D*) and T-type VGCCs (panel *E*). In addition, we investigated the kinetics of the dendritic bAP after blocking the two major types of CAKCs, namely the SK channel using 1 µM apamin (panel *F*) and the BK channel (panel *G*). Consistently with the observation that the blockade of P/Q-type and R-type VGCCs caused, on average, the smallest decrease in the I_Ca_ (Fig.2*C*), in the two representative cells of Fig.6*B* and Fig.6*D* the blockade of these two channels did not change the AP shape. Surprisingly, in the representative cell of Fig.6*E*, the blockade of T-type VGCCs decreased the amplitude of the bAP, suggesting an effect on dendritic excitability. We don’t have an explanation for this effect that can be in principle potentially attributed to the drugs acting on a different target other than T-type VGCC, but this result suggests that the AP weakening may contribute to the relatively large inhibition of the I_Ca_ reported in Fig.2*C*. In all other representative cells, the blockade of the channels caused a visually detectable widening of the AP shape. In the case of N-type VGCCs (Fig.6*C*) and of BK CAKS (Fig.6*G*), a widening seemed occurring during the first 500 µs following the AP peak, therefore during the phase of increasing of VGCC activation. In contrast, in the case of L-type VGCCs (Fig.6*A*) and of SK CAKS (Fig.6*F*), the widening seemed occurring mostly >500 µs after the AP peak, therefore during or after the expected peak of the I_Ca_. The early widening reported in the example of Fig.6*C* was also observed in the example of Fig.7*A*. In this exceptional case, where the peak-to-peak noise was <10 mV, the widening could be visually detected by plotting the difference between the samples after blocking N-type VGCCs and the samples in control conditions. Hence, whereas the 20 differences corresponding to the noise collected before the somatic current injection were either positive or negative values, the 20 differences corresponding to the signal collected after the AP were mostly positive. In the vast majority of tested cells, however, the peak-to-peak noise was >10 mV. Thus, we performed the experiment of the examples of Fig.6 in 8-10 cells for each channel blockade and performed a statistical analysis on the ensemble of the cells tested. Fig.7*B* shows the averaged V_m_ signal of N = 10 cells where the individual traces were aligned with the AP peak. The analysis performed on the 200 sample differences corresponding to the noise indicate that their distribution is consistent with a normal behaviour, according to the large p value obtained by the Lilliefors test. In contrast, the 100 sample differences corresponding to the first 500 µs after the AP peak and to the following 500 µs do not follow a normal distribution. Finally, the hypothesis of deviation from the normal behaviour in the following 1.5 ms could not be rejected. The same analysis was repeated for the blockade of L-type VGCCs (Fig.7*C*), of P/Q-type VGCCs (Fig.7*D*), of R-type VGCCs (Fig.7*E*), of T-type VGCCs (Fig.7*F*), of SK CAKCs (Fig.7*G*) and of BK CAKCs (Fig.7*H*) and the p values obtained by the Lilliefors test are reported in Table 2. A significant deviation from normality in the samples collected in the first 500 µm after the AP peak was discerned in the case of blockade of BK CAKCs. Interestingly, in 4/8 cells tested for the blockade of P/Q-type VGCCs, an early widening of the AP was also observed, leading to a low value of p also in this case. Finally, the significant deviation from normality observed in the case of blockage of T-type VGCCs was due to the consistent reduction of the amplitude of the AP. In contrast to these cases, significant deviations from normality in the samples collected beyond 500 µm after the AP peak were discerned in the case of blockade of L-type VGCCs and in the case of blockade of SK CAKCs. These behaviours were due to the widening of the AP shape during or after the expected occurrence of the I_Ca_ peak. In the case of these two blockades, the time-course of the AP change is consistent with an activation of K^+^ channels by cytosolic Ca^2+^, but the hypothesis that these channels are also selectively activated by the specific Ca^2+^ source cannot be excluded since the delayed widening can be due to a slower kinetics of the K^+^ channels. Yet, in the case of the blockade of N-type VGCCs or of the blockade of BK CAKCs, the results indicate that the activation of K^+^ channels is not linear with the cytosolic Ca^2+^, preceding the peak of the I_Ca_. The fact that the AP widening occurs when only a fraction of inflowing Ca^2+^ via N-type VGCCs is detected indicates that BK channels are activated before the Ca^2+^-dye binding reaction equilibrates. This result suggests a physical interaction between the N-type VGCC and the BK CAKC at nanoscopic domain. Thus, we further explored this hypothesis using biophysical modelling in the NEURON environment.

**Figure 6.**
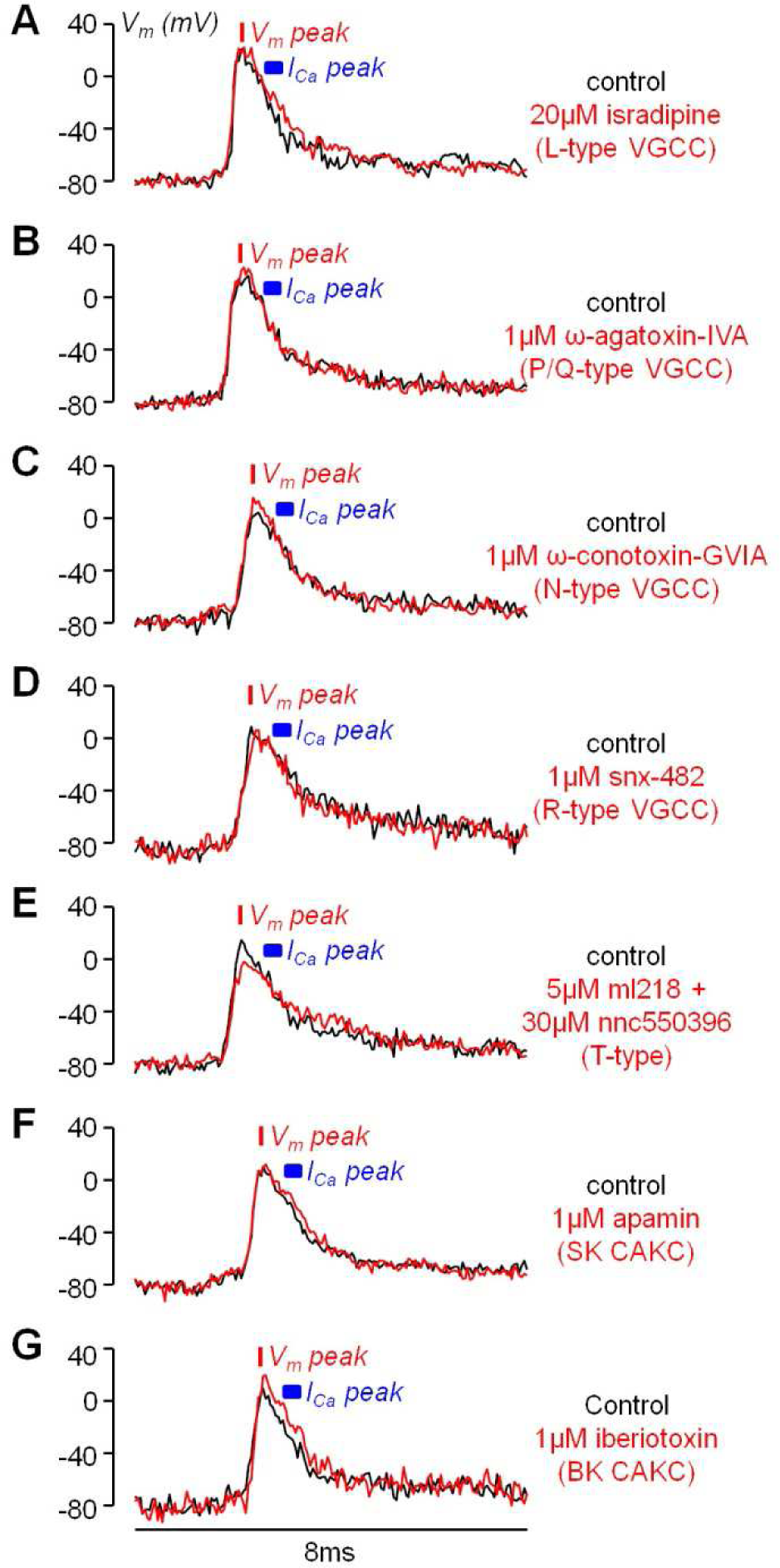
Examples of the effects of the diverse VGCC and CAKC blockades on the bAP shape. ***A***, Optical V_m_ transient from an apical dendrite region (∼100 µm from the soma) in a representative L5 pyramidal neuron in control conditions (black trace) and after blocking L-type VGCCs with 20 µM isradipine (red trace). ***B***, Representative experiment as in *A*, but by blocking P/Q-type VGCCs with 1 µM ω-agatoxin-IVA. ***C***, Representative experiment as in *A*, but by blocking N-type VGCCs with 1 µM ω-conotoxin-GVIA. ***D***, Representative experiment as in *A*, but by blocking R-type VGCCs with 1 µM snx-482. ***E***, Representative experiment as in *A*, but by blocking T-type VGCCs with 5 µM ml218 and 30 µM nnc5503961. ***F***, Representative experiment as in *A*, but by blocking SK CAKCs with 1 µM apamin. ***G***, Representative experiment as in *A*, but by blocking BK CAKCs with 1 µM iberiotoxin. In each cell the red bar shows the timing of the AP peak and the blue segment the expected range for the peak of the I_Ca_ (500-800 µs after the AP peak).

**Figure 7.**
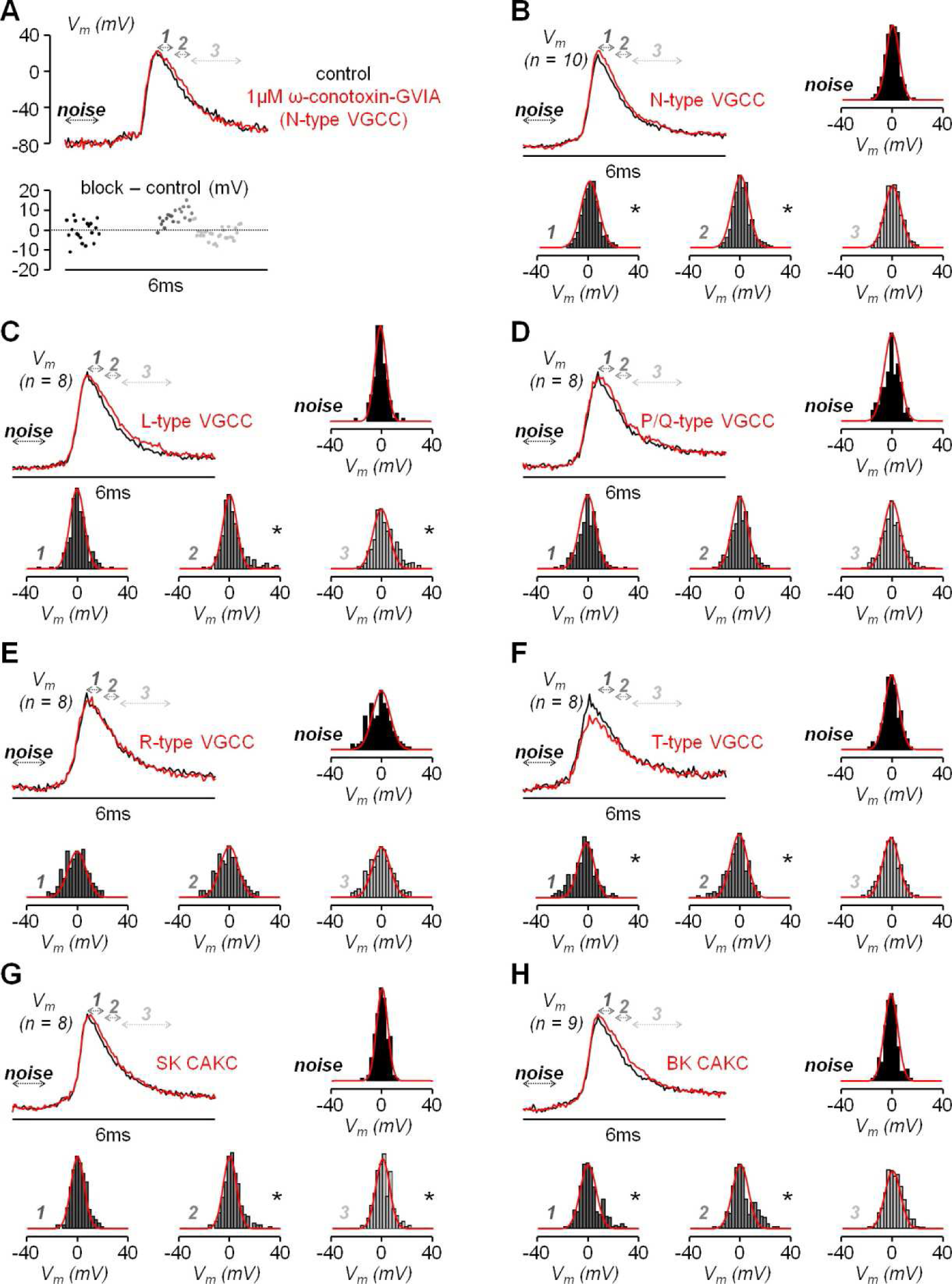
Analysis of the effect of the diverse VGCCs and CAKs on the bAP shape. ***A***, Top, optical V_m_ transient from an apical dendrite region (∼100 µm from the soma) in a representative L5 pyramidal neuron in control conditions (black trace) and after blocking N-type VGCCs (red trace); arrows at different gray tones indicate the time windows corresponding to the noise (1 ms before current injection), signal “*1*” (first 500 µs after the AP peak), “*2*” (following 500 µs) and “*3*” (following 1.5 ms). Bottom, sample difference of the signals after the channel blockade and in control conditions in the four time windows. ***B***, Averaged signals aligned to the AP peak from N = 10 cells in control conditions (black trace) and after blocking N-type VGCCs (red trace) and histograms of the sample differences in the illustrated four time windows; normal distributions with mean and standard deviations calculated from the points are reported (red plots) for comparison with the histograms. ***C***, Same as in *B*, but in 8 cells where L-type VGCCs were blocked. ***D***, Same as in *B*, but in 8 cells where P/Q-type VGCCs were blocked. ***E***, Same as in *B*, but in 8 cells where R-type VGCCs were blocked. ***F***, Same as in *B*, but in 8 cells where T-type VGCCs were blocked. ***G*** Same as in *B*, but in 8 cells where SK CAKs were blocked. ***H***, Same as in *B*, but in 9 cells where BK CAKs were blocked. In panels *B-H* “*” indicates that the distribution deviates from normality (p < 0.01, Lilliefors test).

**Table 2.**
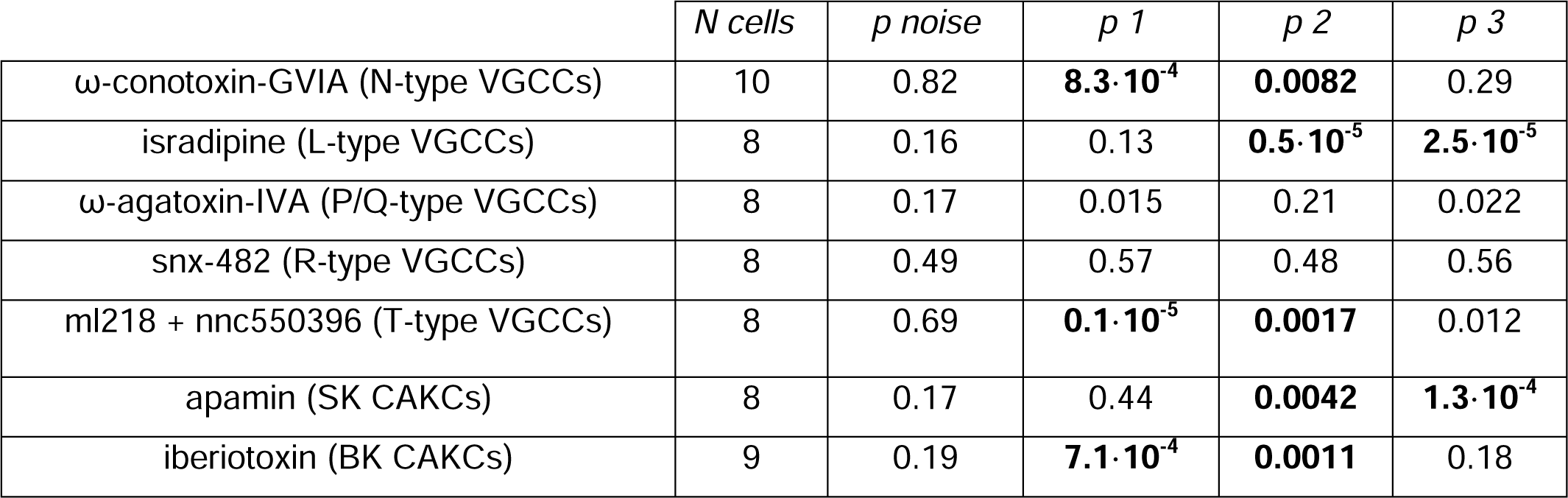
Lilliefors test to assess whether a set of values is consistent with a normal distribution performed on sample differences in V_m_ imaging experiments after blocking a channel (with the name of the blocker indicated) and in control conditions. The five columns report, for each channel tested, the total number of cells (*N*) and the values of p (*p*) for the noise (number of samples 20·*N*), for the signal *1* (first 500 µs after the AP peak, number of samples 10·*N*), for the signal *2* (following 500 µs after the AP peak, number of samples 10·*N*), for the signal *3* (following 1.5 ms after the AP peak, number of samples 30·*N*). Tests where p was < 0.01 are reported in bold characters and indicate a significant change of the AP shape caused by the blockade of the channel.

### The coupling between N-type VGCCs and BK CAKCs is confirmed by a NEURON model

In order to reproduce the kinetics of the AP and of the I_Ca_, we built a realistic NEURON model, and considered an apical dendrite at ∼100 µm from the soma, corresponding to the zone explored in the experiments. The model included all types of VGCCs tested, BK CAKCs and SK CAKCs. The coupling between a specific Ca^2+^ channel and the K^+^ channel was modelled by introducing an activation factor (α) of the BK channel by the Ca^2+^ influx from that particular channel. This coupling was imposed both to L-type and N-type VGCCs, in agreement to what reported in the literature (Sun et al., 2003). We found that, to replicate the ensemble of experimental results, the coupling needed to be strong for N-type VGCCs (α = 10) and weak for N-type VGCCs (α = 3). Fig. 8*B* shows the dendritic AP, which reproduces the experimental AP obtained in one cell, superimposed to the Ca^2+^ transient and of the I_Ca_. Consistently with experimental observations, the model showed a delay of ∼0.8 ms between the peak of the AP and the peak of the I_Ca_. Next, to mimic the pharmacological blockade of channels, from the model corresponding to the control condition we removed either 90% of N-type VGCCs, 90% of BK CAKCs or 90% of both channels (Fig. 8*B*). The model reproduced the experimental widening of the AP preceding the peak of the I_Ca_ and the increase in the Ca^2+^ signals with all simulated pharmacological blockades. In addition, consistently with the results of the experiments reported in Fig.4, the blockade of both channels decreases the Ca^2+^ signals with respect to the simulation where only BK CAKSs are blocked. To assess whether the selective coupling between N-type VGCCs and BK CAKAs was necessary to reproduce the experimental results, we suppressed the coupling between N-type VGCCs and BK CAKCs. In this modified model, the early widening of the AP disappears and the Ca^2+^ signal and current decrease when N-type VGCCs are blocked (Fig. 8*C*). This result supports the experimental suggestion that a strong coupling between N-type VGCCs and BK CAKAs may be in effect in these neurons.

**Figure 8.**
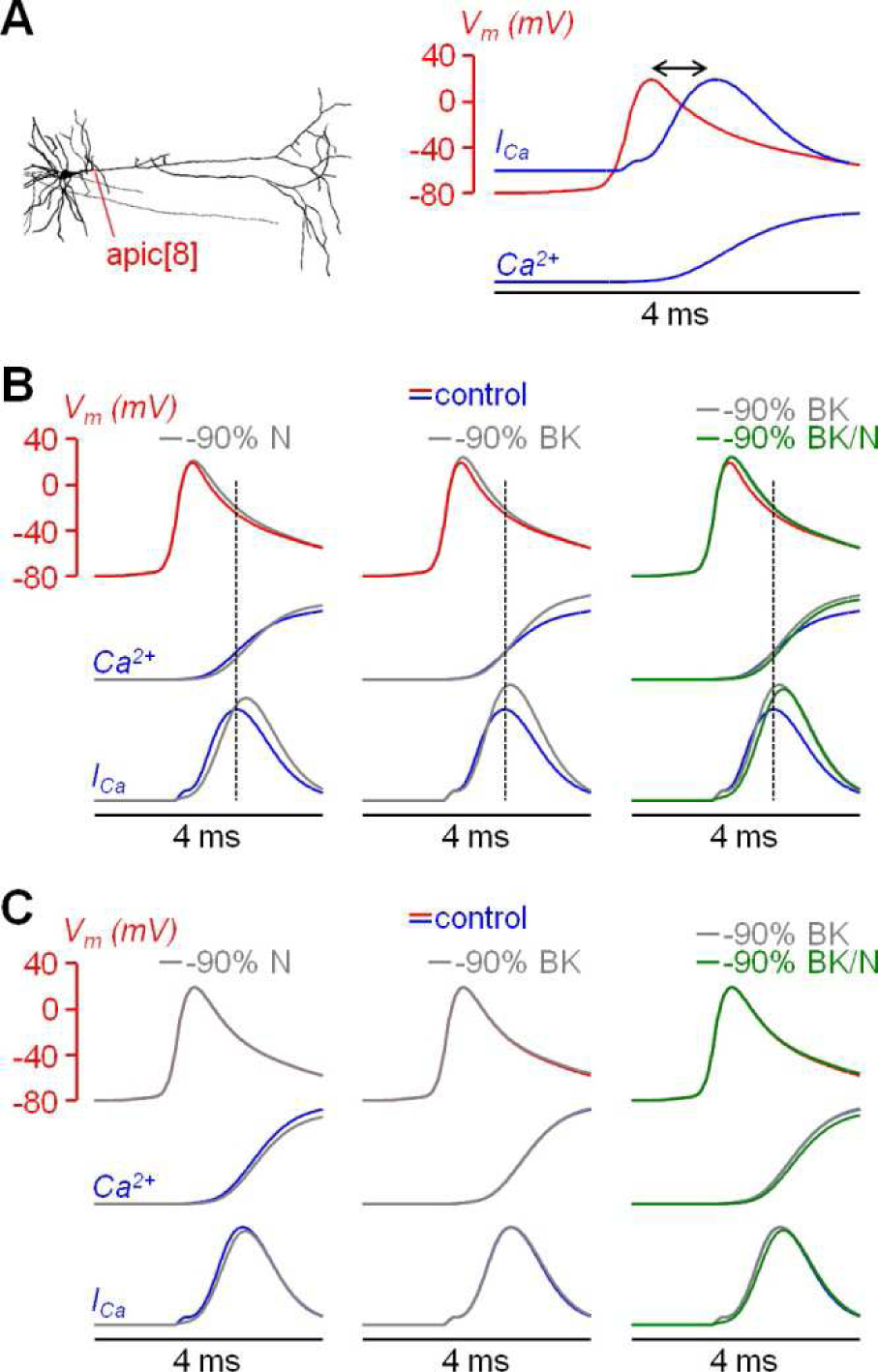
Effects of coupling between BK CAKCs and N-type VGCCs in a biophysically accurate model. ***A***, *Left,* the morphology of the L5 pyramidal neuron used in all simulations; the apical dendrite used to compare model and experiments is indicated in red. *Right*, dendritic AP evoked by a somatic current injection (red trace); time course of the associated Ca^2+^ transient and current (blue traces); the delay between the peaks of the AP and the I_Ca_ is indicated by the double arrow. ***B***, dendritic AP under control conditions (red trace) and the kinetics of the associated Ca^2+^ transient and current (blue traces); gray traces on the left are after removing 90% of N-type VGCCs; gray traces in the middle and right are after removing 90% of BK-type CAKCs; green traces on the right are after removing 90% of N-type VGCCs and BK-type CAKCs. Dotted lines indicate the timing of the I_Ca_ peak. Note the agreement with experimental recordings. ***C***, same as in panel B, but from simulations in a model with no coupling between N-type VGCCs and BK CAKCs. Note that traces are not consistent with experimental results.

## DISCUSSION

In this report we provide strong experimental evidence that BK CAKCs are selectively activated by N-type VGCCs in the apical dendrite of L5 pyramidal neurons. This activation is rapid and is not linear with the increase of intracellular Ca^2+^ concentration associated with the transient activation of VGCCs. Thus, the blockade of N-type VGCCs widens the AP peak prolonging Ca^2+^ entry through the other VGCCs and boosting the increase of intracellular Ca^2+^ concentration associated with the AP. This result also indicates that our imaging approach can indirectly reveal functional protein-protein interactions that can be foreseen using structural imaging techniques such as Förster resonance energy transfer imaging (Masi et al., 2010). Indeed, in the case of VGCC nanodomains (Gandini and Zamponi, 2021), the close interaction with the Ca^2+^ target enables sub-millisecond activation of the target, as has been shown in synaptic terminals (Volynski and Krishnakumar, 2018). Here, we show that the combined analysis of cytosolic Ca^2+^ and of AP kinetics at 50 µs temporal resolution can reveal VGCC-CAKC interactions at functional level in larger structures such as neuronal dendrites.

### Functional consequences of the selective coupling between BK and a specific calcium source

Overall, BK channels are known to perform multiple functions in neurons (Ancatén-González et al., 2023). Compared with other CAKCs, BK channels exhibit lower affinity to Ca^2+^ and require concomitant depolarisation (Berkefeld et al., 2010). These two biophysical properties suggest that these channels must be physically close to a specific Ca^2+^ source and that they can activate at a sub-millisecond time scale when the cell is depolarised (Berkefeld et al., 2006). It is known that BK CAKCs form macromolecular complexes with L-type, P/Q-type and N-type VGCCs and, when the two proteins were co-expressed in heterologous systems, it was found that the kinetics of activation of BK CAKCs depended on the coupled VGCCs (Berkfeld and Fakler, 2008). In presynaptic terminals, BK channel activity regulate synaptic transmission (Raffaelli et al., 2004) and can modulate both the amplitude and duration of depolarization-evoked Ca^2+^ entry as a result of the rapid repolarization and deactivation of P/Q-type and N-type VGCCs (Fakler and Adelman 2008). In turn, reduced Ca^2+^ influx limits vesicle fusion at active zones, leading to decreased neurotransmitter release (Kyle and Braun 2014). In the postsynaptic areas of L5 pyramidal neurons, BK channels are expressed not only in the dendritic bulk, but also in synaptic spines, and activation of BK CAKCs in small-head spines by Ca^2+^ influx through glutamate receptors reduces the size of synaptic potentials (Tazerart et al. 2022). Thus, both in synaptic terminals and spines, the selective activation of BK channels by a specific partner leads to a negative feedback of the triggering signal, which is the AP in the case of synaptic terminal and the synaptic potential in the case of spines. This negative feedback translates into a reduction of either neurotransmitter release or of neuronal excitability once the synaptic potential has reached the soma and the axon initial segment. Thus, in both cases, the functional consequences of BK CAKC activation can be tied to the underlying K^+^ current that decreases the amplitude and/or the size of the triggering depolarizing event. In the apical dendrite of L5 pyramidal neurons, we found that BK CAKCs activation by N-type VGCCs anticipates the full activation of VGCCs, providing a negative feedback to the Ca^2+^ current also mediated by other channels. The AP shaping produced by BK CAKCs is qualitatively similar to what we observed in the axon initial segment of L5 pyramidal neurons, where these channels are activated by the Ca^2+^ permeable voltage-gated Na^+^ channel Na_v_1.2 (Filipis et al., 2023). Compared to synaptic terminals and spines, however, the role of the regulation of the AP shape in these wider regions, produced by the K^+^ current, is less straightforward.

### Putative functional consequences of BK membrane conductance increase

Ion channels do not only mediate ionic currents that change the V_m_, but they can also regulate the spread of V_m_ transient from a site to another by locally changing the membrane conductance, i.e. by “shunting” the transmission of the signal when the channels open (Blomfield, 1974). Whereas shunting inhibition is traditionally associated with localised increases of membrane conductance, the AP represents a very brief increment of membrane conductance that propagates along dendritic branches. Our realistic model shows that BK CAKCs represent a significant fraction of this membrane conductance transient that is timely-locked to the AP. Thus, it is possible that the BK conductance transient can effectively contribute to the firing modulation produced by incoming synaptic inputs which are briefly shunted by the back-propagating AP. As for the case of localised BK channel activation in synaptic terminals and spines, the functional consequence of the spread BK channel activation is a reduction of neuronal excitability, in this case occurring at the level of dendritic integration.

### Potential relevance in neurological disorders

The N-type VGCC, which is expressed in neurons predominantly at presynaptic terminals, is associated with several neurological conditions such as anxiety, addiction, and pain in correlation with its endogenous regulator nociceptin opioid peptide receptor (Caminski et al. 2022). In contrast, dysfunction of BK CAKCs encoded by the *KCNMA1* gene, which are widely expressed in many tissues, is associated with complex combinations of disorders, including seizures, movement disorders, developmental delay and intellectual disability (Bailey et al. 2019). For instance, a *KCNMA1* knock-out mouse exhibits motor impairments and suffers from learning difficulties (Typlt et al. 2013), whereas paroxysmal non-kinesigenic dyskinesia is observed in patients with gain-of-function channelopathies of BK channels (Miller et al. 2021). Notably, BK channel dysfunction is also reported in other genetic diseases such as the fragile X syndrome (Deng and Klyachko, 2016), suggesting that BK CAKCs can be a potential general target for therapeutic intervention (Griguoli et al., 2016). In the case of the signal described in the present report, the specific coupling of BK CAKCs to N-type VGCCs in the apical dendrite might be determined by alternative splicing of the channel or through auxiliary subunits. It is known that BK CAKCs are regulated by extensive alternative splicing as well as multiple auxiliary subunits, giving BK CAKCs both cell and tissue-specific properties (Kyle and Braun 2014; Latorre et al. 2017), even within the L5 pyramidal neuron population (Guan et al. 2015). In recent years, various β and γ auxiliary subunits of BK channels have been identified which can modulate activation and inactivation dependencies (Gonzalez-Perez and Lingle 2019), but the specific co-expression can in principle also drive the choice of the Ca^2+^ channel partner in sub-cellular compartments. We have previously reported that in the axon initial segment, BK CAKCs interact with Na_v_1.2 channels, but this interaction seems independent of auxiliary subunits as was indicated by heterologous expression of these channels in HEK293 cells (Filipis et al. 2023). In contrast, the BK CAKCs could be guided by auxiliary subunits to form selective nanodomains with N-type VGCCs in the dendrites of L5 pyramidal neuron. This hypothesis requiring extensive investigation might be potentially important to investigate the specific function of BK CAKC characterised in this report.

## Acknowledgements

This work was supported by the *Agence Nationale de la Recherche* through two grants (ANR-18-CE19-0024 – OptChemCom and Labex *Ion Channels Science and Therapeutics:* program number ANR-11-LABX-0015). E.G. was funded by the Regione Sicilia (CUP G79J21012770001). We are indebted to the *Fédération pour la Recherche sur le Cerveau* (FRC – Grant *Espoir en tête*, Rotary France) and the National Infrastructure France Life Imaging for financing part of the experimental equipment. We acknowledge a contribution from the Italian National Recovery and Resilience Plan (NRRP), M4C2, funded by the European Union – NextGenerationEU (Project IR0000011, CUP B51E22000150006, “EBRAINS-Italy”).

## Notes

https://doi.org/10.5281/zenodo.7623898

### Competing Interest Statement

The authors have declared no competing interest.

